# Association mapping identified novel candidate loci affecting wood formation in Norway spruce

**DOI:** 10.1101/292847

**Authors:** John Baison, Amarylis Vidalis, Linghua Zhou, Zhi-Qiang Chen, Zitong Li, Mikko J. Sillanpää, Carolina Bernhardsson, Douglas Scoffield, Nils Forsberg, Lars Olsson, Bo Karlsson, Harry Wu, Pär K. Ingvarsson, Sven-Olof Lundqvist, Totte Niittylä, M Rosario García-Gil

## Abstract

➢ Norway spruce (*Picea abies*) is an important boreal forest tree species of significant ecological and economic importance. Hence there is a strong imperative to dissect the genetics controlling important wood quality traits in the species.
➢ We performed a functional genome-wide association mapping of 17 wood traits in Norway spruce using 178101 single-nucleotide polymorphisms (SNPs) generated from exome genotyping of 517 mother trees. The wood traits were defined using functional modelling of wood properties across annual growth rings.
➢ Association mapping was performed using a multilocus LASSO penalized regression method and we detected a total of 51 significant SNPs from 39 candidate genes that are involved in wood formation.
➢ Our study represents the first functional multi-locus genome-wide association mapping (AM) in Norway spruce. The results advance our understanding of the genetics influencing wood traits, identify novel candidate genes for further functional studies and support current Norway spruce breeding efforts.

## Introduction

Norway spruce (*Picea abies* (L.) Karst.) is a dominant boreal softwood species of significant economic and ecological importance (Hannrup *et al*., 2004). Long-term Norway spruce breeding programmes for improvement of growth and survival were initiated in the 1940s and recently, wood quality has become one of the priority traits (Bertaud & Holmbom, 2004; Hannrup *et al*., 2004). Norway spruce breeding in Sweden complete one cycle in about 20 years and such long generation times make improvements in growth and wood quality very slow. Among wood quality traits, wood density is considered a key indicator of stability, strength and stiffness of sawn timber (Hauksson *et al*., 2001). Several studies of wood quality observed that fast growth conflicts with high quality wood, as shown by the negative genetic correlation between wood volume growth and density in Norway spruce (Olesen, 1977; Dutilleul *et al*., 1998; Chen *et al*., 2014). In order to combine fast growth and desirable wood properties through breeding, and to shorten the breeding cycle, it is therefore imperative to design effective early selection methods and breeding strategies. In an effort to design optimal breeding and selection strategies for reducing or breaking negative genetic correlations between traits it is essential to identify alleles that are responsible for generating favourable or unfavourable genetic correlations (Hallingbäck *et al*., 2014).

When DNA markers were first introduced in 1980s, tree breeders were provided a possibility to correlate phenotypes with polymorphic DNA markers and to conduct selection using genotypes instead of phenotypes (Lande & Thompson, 1990). Groover et al. (1994) first identified quantitative trait loci (QTL) for wood density variation in loblolly pine using linkage analyses based on segregating family pedigrees. However, maker-aided selection (MAS) based on results from QTL analyses was never implemented in practical tree breeding due to the so-called Beavis effect (e.g. inflated estimates of allelic effects and underestimation of QTL number for economically important traits) (Beavis, 1998), inconsistent associations among different families and the low transferability of markers (Strauss *et al*., 1992). Association Mapping (AM) is a more powerful QTL detection method that was introduced to tree genetics using a candidate gene approach (Thumma *et al*., 2010). AM overcomes the limited resolution of family-based QTL mapping by relying on historical recombination in the mapping population (Neale & Savolainen, 2004; Thavamanikumar *et al*., 2013; Huang & Han, 2014). The effectiveness of AM relies on genome-wide levels of LD, which decays rapidly within coding regions in conifer species, however, it may be extensive in certain non-coding regions (Moritsuka *et al*., 2012). Fast-decaying LD, coupled with complex polygenic nature for both growth and wood quality traits (Hall *et al*., 2016) implies that a large number of genomic regions need to be investigated to identify significant QTL (Beaulieu *et al*., 2011).

The availability of a draft genome sequence for Norway spruce (Nystedt *et al*., 2013) has opened new possibilities for the development of genetic markers to conduct both AM at the genome-wide level (genome-wide association, GWAS) and genomic selection (GS). Several reduced representation-based approaches such as sequence capture and transcriptome sequencing (Hirsch *et al*., 2014) have been developed as complexity-reduction methods suited for studying large genomes, such as the 20Gb Norway spruce genome. These approaches reduce the sequence space by decreasing the repetitive sequence content of the genome. In this study we employed a solution-based sequence capture method.

Several AM studies have been performed in trees and have identified genetic loci linked to, for instance, wood properties in *Populus trichocarpa (Porth et al., 2013)*, adaptive traits in *Pinus contorta* (Parchman *et al*., 2012) and to wood quality traits in *Eucalyptus* (Porth *et al*., 2013; Resende *et al*., 2012). Such studies aimed at dissecting the genetic basis of wood properties can benefit from the application of mathematical functions that account for year-to-year variation across annual growth rings, cambial age and distance from pith (Li *et al*., 2014). Mathematical modelling allows the incorporation of phenotypic growth trends that increase the precision and resolution of QTL detection through the integration of the phenotype information over multiple time points and reduction of residual variance (Ma *et al*., 2002). Such functional mapping analysis can be conducted using a multistage approach (Heuven & Janss, 2010). First, the phenotype trends of each individual are modelled using curve-fitting methods and the parameters describing the curve are then considered as latent traits. The latent traits are then used in an independent association analyses to search for genomic regions affecting the trait and to estimate genetic marker effects (Li *et al*., 2014).

In this study, we applied a functional genome-wide association mapping (AM) approach to identify genomic regions contributing to wood quality traits in Norway spruce [*Picea abies* (L.) Karst.]. Estimated breeding values (EBVs) were calculated for growth and wood quality traits at the resolution of annual growth rings and were then used to extract latent traits from fitting quadratic splines, Fig. 1a. We applied quadratic splines since traditional analyses that utilise a single point data across annual growth rings may confound the analyses by averaging across a full sample. Such averaging may obscure mechanisms acting at specific time points during wood formation and will make identification of underlying genes more difficult. In this study, we have refined our data in order to cover within ring features in earlywood (EW) and latewood (LW), as well as in a more weather influenced part in between named transitionwood (TW). This study has also performed the first analysis of number of cells per ring calculated from SilviScan data. Penalized LASSO regression (Tibshirani, 1996) and the stabilizing selection probability method of (Meinshausen & Bühlmann, 2010) were then used, Fig. 1c, to detect significant associations between latent traits derived from EBVs and 178101 SNP markers covering the Norway spruce genome, Fig. 1b.

**Fig 1.**
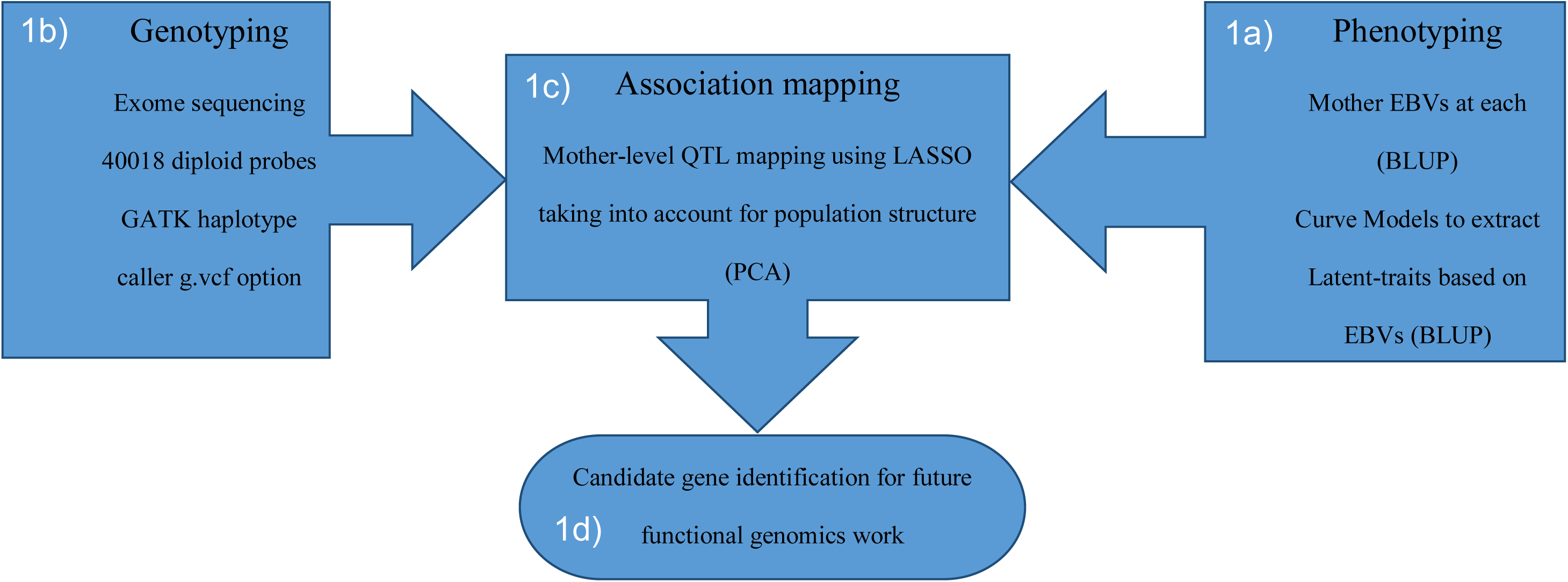
Outline of the association mapping approach: 1a) Mother Estimated breeding values (EBVs) were determined using a univariate, bivariate or multivariate mixed linear model based on the different fitness of the model with the resultant values adjusted with the mean. The adjusted EBVs were plotted against cambial age (annual ring number) to produce time trajectories for each trait. A quadratic spline curve model was then applied to the EBVs to estimate latent-traits. 1b) Sequence capture on the 517 from 40018 diploid probes resulted in 178101 single-nucleotide polymorphisms (SNPs). 1c) Association mapping was the performed using a multi-locus LASSO penalized regression method with PCA components acting as covariates accounting for the population structure in our mother trees. 1d) Finally, a candidate gene identification process for contigs with significant SNPs was conducted in ConGenIE and public sequence databases.

## Materials and Methods

### Plant material and phenotype data

Plant material and phenotype data used in this study have previously been described in Chen et al. (2014). In brief, two progeny trails were established in 1990 in Southern Sweden (S21F9021146 aka F1146 (trial1) and S21F9021147 aka F1147 (trial2)). These trials were composed of 1373 and 1375 open pollinated families, respectively, and form the basis of our analyses. We selected 517 families in 112 sampling stands to use in the investigation of wood properties. At each site, increment wood cores of 12 mm were collected from six trees of the selected families at breast height (1.3 m) (6 progeny × 2 sites = 12 progenies in total). A total of 5618 trees, 2973 and 2645 trees from the F1146 and F1147 trails respectively, were analysed. The pith to bark profiling of the wood physical attributes was analysed using the SilviScan technology (Evans 1994, 2006) at Innventia, Stockholm, Sweden, where the initial data evaluations were performed using customized methods. These methods focus on the identification and dating of all annual rings and their compartments of earlywood, transitionwood and latewood. For the current study, Innventia also calculated three additional traits, number of cells per ring (NC), wood percentage (WP), and a trait named Mass Index (MI), introduced to express the relative amount of biomass, all derived from the SilviScan data. MI was then used to identify trees with an uncommon positive correlation between density and growth, that is more biomass. The traits included in the current study are listed in Table 1.

**Table 1.**
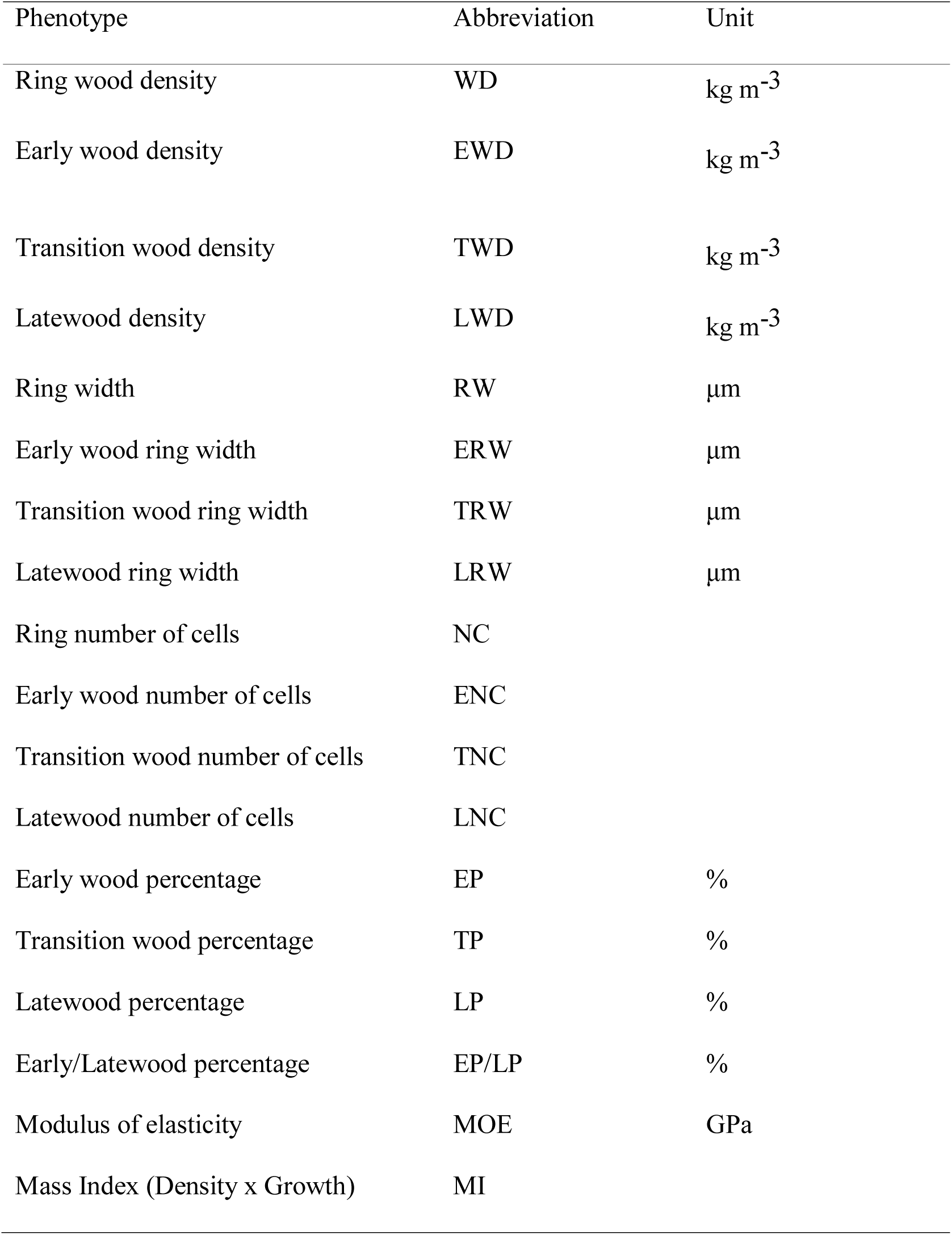
List of the phenotypes, their abbreviations and measurement unit.

| Phenotype | Abbreviation | Unit |
| --- | --- | --- |
| Ring wood density | WD | kg m<sup>-3</sup> |
| Early wood density | EWD | kg m<sup>-3</sup> |
| Transition wood density | TWD | kg m<sup>-3</sup> |
| Latewood density | LWD | kg m<sup>-3</sup> |
| Ring width | RW | μm |
| Early wood ring width | ERW | μm |
| Transition wood ring width | TRW | μm |
| Latewood ring width | LRW | μm |
| Ring number of cells | NC |  |
| Early wood number of cells | ENC |  |
| Transition wood number of cells | TNC |  |
| Latewood number of cells | LNC |  |
| Early wood percentage | EP | % |
| Transition wood percentage | TP | % |
| Latewood percentage | LP | % |
| Early/Latewood percentage | EP/LP | % |
| Modulus of elasticity | MOE | GPa |
| Mass Index (Density x Growth) | MI |  |

### Statistical analysis

EBVs were calculated for each cambial age (annual ring) separately and used for statistical modelling to derive latent traits. The variance and covariance components were estimated using ASREML 4.0 (Gilmour *et al*., 2014) as described in Chen et al., (2014). In brief, the EBVs at each cambial age were estimated using a univariate, bivariate or multivariate mixed linear models. The fit of different models were evaluated using the Akaike Information Criteria (AIC) and the optimal model was selected based on a compromise of model fit and complexity. Breeding values were then centred in order to obtain within genotype trends. A univariate linear mixed model for joint-site analysis was implemented as:

The following univariate linear mixed model for joint-site analysis was fitted as:

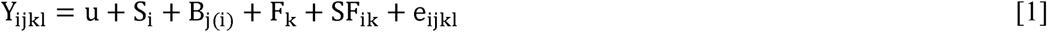
where Y_ijk1_ is the observation on the *l*th tree from the *k*th family in jth block within the *i*th site, *u* is the general mean, S_i_ and B_j(i)_ are the fixed effects of the *i*th site and the *j*th block within the ith site, respectively, F_k_ and SF_ik_ are the random effects of the *k*th family and the random interactive effect of the *i*th site and *k*th family, respectively, e_ijk1_ is the random residual effect. Multivariate mixed linear models were used to estimate BV for different phenotype traits if the model fitted better than bivariate or univariate based on AIC.

A number of trees were observed that broke the negative correlation usually observed between density and growth. These trees exhibited both high density and fast growth, thus larger biomass. In order to identify putative genes involved in this favourable combination of traits, we defined a new trait termed Mass Index (MI), that we subsequently used in the association mapping. The MI was defined as follows:

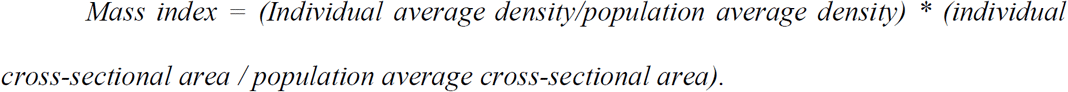

The index was then treated as a new dynamic trait in the AM analyses where individuals with an index > 1 indicate a wood mass per length unit than the population average in the cross-section at breast height. The index was calculated for all progeny and were used to calculate BVs for the 517 mother trees.

The EBVs were plotted against cambial age (annual ring number) to produce time trajectories for each trait (Fig. 2 and Fig. S1) and used to estimate latent curve parameters. At the first stage, all the trajectories versus cambial age were fitted with a quadratic spline with multiple knots in order to describe the dynamics of the EBVs across age. In this study, this was done with the values of four parameters obtained from the spline fitting: the intercept, the slope and two knot parameters (K1 and K2). The intercept and slopes were used to evaluate the mean and rate of change for the trait across the annual rings, respectively. K1 and K2 represent inflection points in the cambial age trajectories where the development of the EBVs enters new phases. These two points (K1 and K2) are therefore supposed to have biological significance, warranting a closer analysis of the genes imparting these shifts in the EBVs dynamics. The four latent traits show lower correlations compared to the direct measurements on the original scales and they also have constant variances, thereby reducing the need to account for residual dependencies in the model (Li *et al*., 2014).

**Fig 2.**
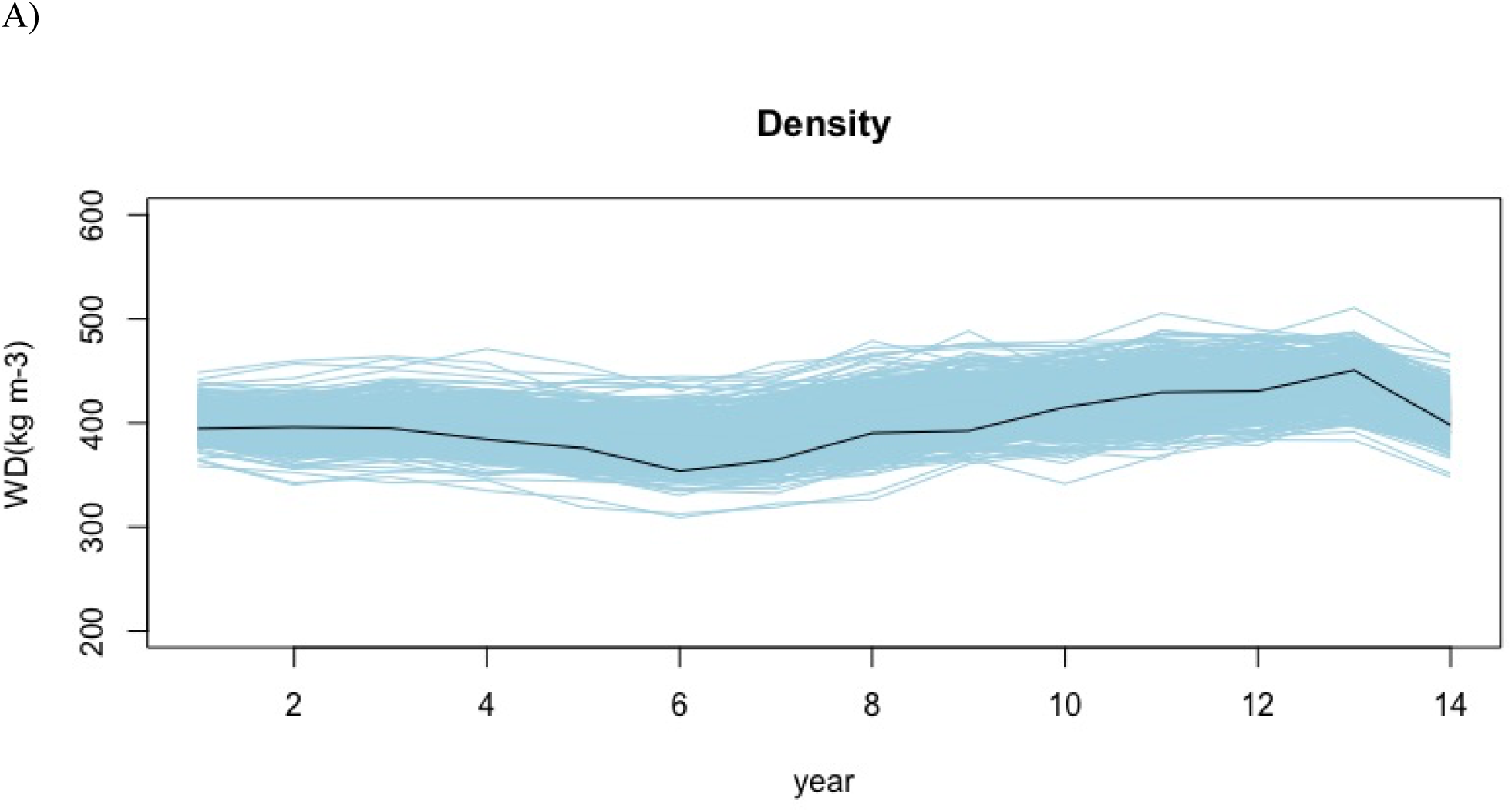
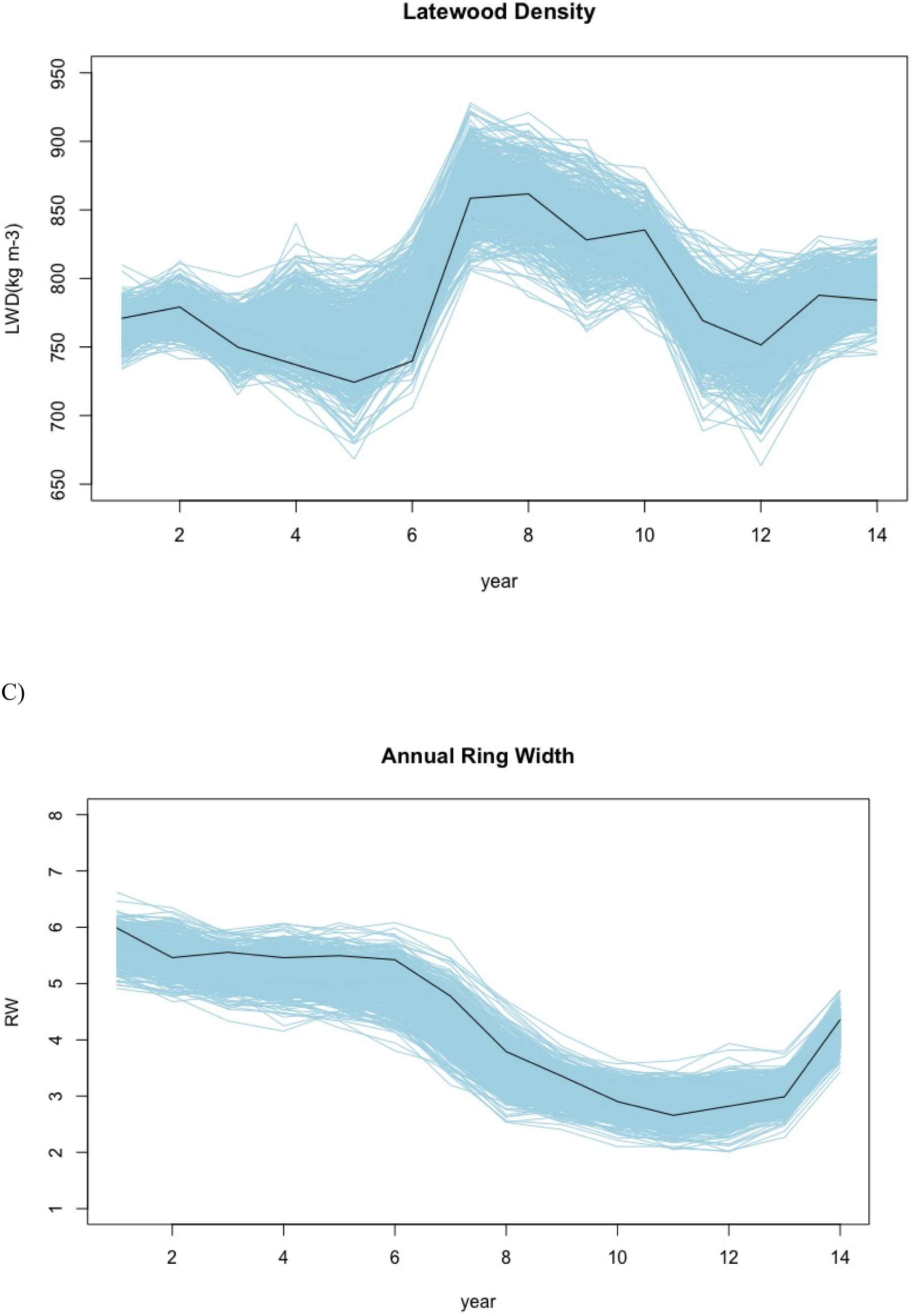
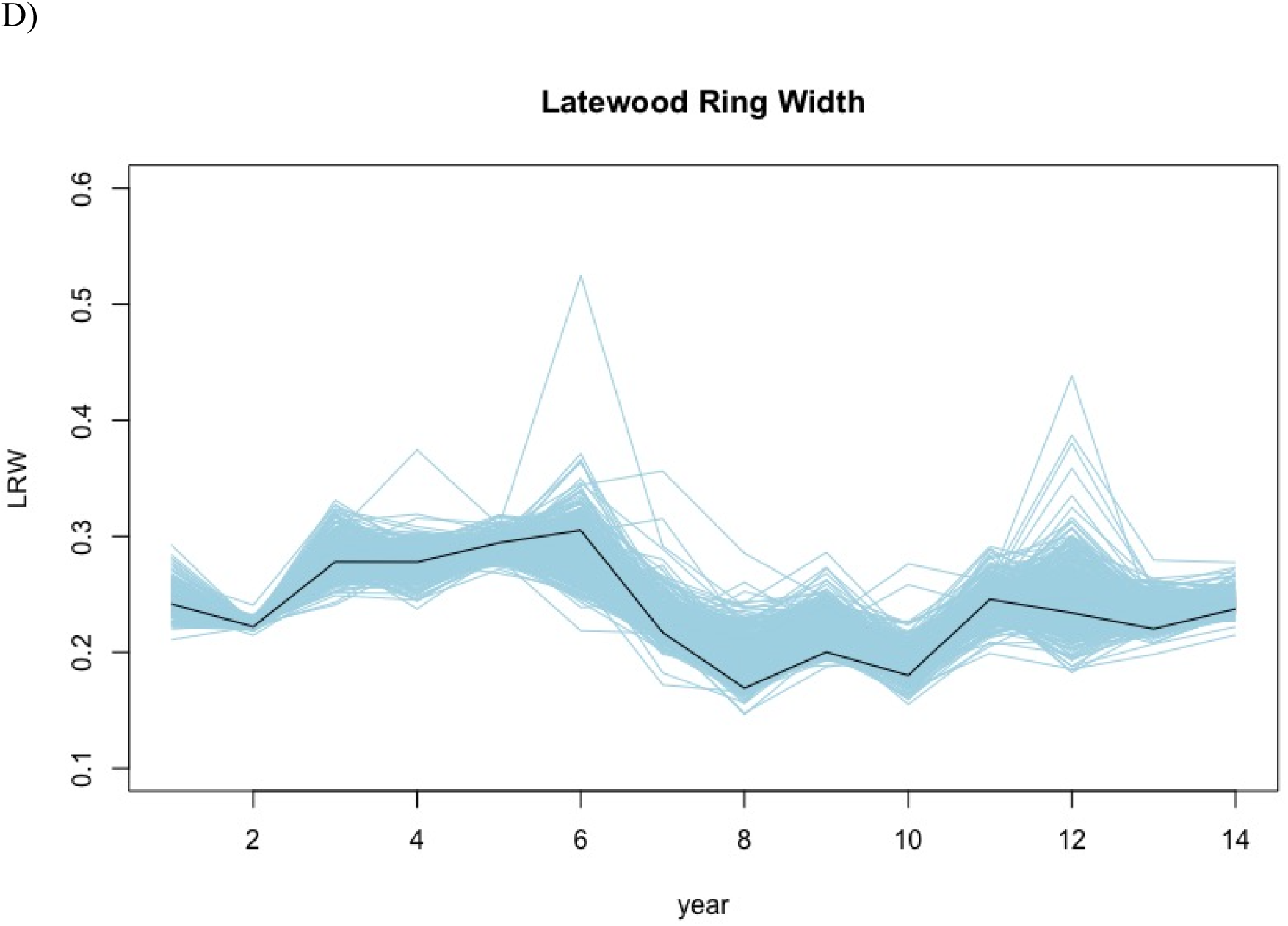
EBV trajectories of four wood quality traits by time:(A) wood density, (B) latewood density, (C) annual ring width and (D) latewood ring width. Individual trajectories for each trait are shown in light blue lines and the black line represents the mean trajectory for the phenotype. These individual trajectories were used to determine the four latent traits of each tree, using quadratic splines with two knots.

The general definition of a quadratic spline with multiple knots is as follows:

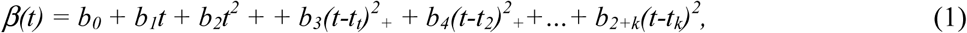
which is continuous and where *t*_i_ (*i*=1,…,*k*; *t*_1_<*t*_2_…<*t_k_*) are defined as knots, and *(t − t_i_)^2^+ = (t − t_i_)^2^* if *t > t_i_* (*t_i_*>0; *i*=1,…,*k*), and otherwise is equal to zero. The number of knots has to be properly defined in order to provide an accurate description of the data under investigation, as well as functional starting points for the search of their locations (Li *et al*., 2015). In our case, since the growth pattern of wood property traits were not complex, we choose two knots of the time interval.

Hence, the quadratic spline model to describe the growth trajectory of individual *i* applied in this study was defined as:

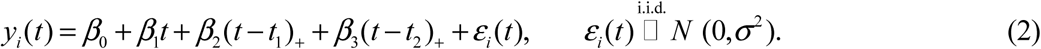

Then the intercept *β*_0_, slope *β*_1_, *β*_2_, (Knot 1 (k1)) and *β*_3_ (Knot 2 (k2)) are estimated by standard least squares, and their estimates were considered as the latent trait in the subsequent QTL analysis conducted in R-studio (Team, 2015). The latent traits were then analysed using the LASSO model in order to identify SNPs showing significant associations to the traits.

### Sequence capture, genotyping and SNP annotation

Total genomic DNA was extracted from 517 unrelated individuals using the Qiagen Plant DNA extraction protocol with DNA quantification performed using the Qubit® ds DNA Broad Range (BR) Assay Kit (Oregon, USA). Sequence capture was performed using the 40 018 diploid probes previously designed and evaluated for *P. abies* (Vidalis *et al*., 2018) and samples were sequenced to an average depth of 15x using an Illumina HiSeq 2500 (San Diego, USA). Raw reads were mapped against the *P.abies* reference genome v1.0 using BWA-mem (Langmead & Salzberg, 2012; Li & Durbin, 2009). SAMTools v.1.2 (Li *et al*., 2009) and Picard v.1.140 (McKenna *et al*., 2010) were used for sorting and removal of PCR duplicates. Variant calling was performed using GATK HaplotypeCaller v.3.6 (McKenna *et al*., 2010) in gVCF output format. Samples were then merged into batches of ~200 before all 517 samples were jointly called.

Variant Quality Score Recalibration (VQSR) method was performed in order to avoid the use of hard filtering for exome/sequence capture data. For the VQSR analysis two datasets were created, a training subset and input file. The training dataset was derived from a Norway spruce genetic mapping population with loci showing expected segregation patterns (Bernhardsson *et al*., 2018) and assigned a prior value of 15.0. The input file was derived from the raw sequence data using GATK best practices with the following parameters: extended probe coordinates by +100 excluding INDELS, excluding LowQual sites, and keeping only bi-allelic sites. The following annotation parameters QualByDepth (QD), MappingQuality (MQ) and BaseQRankSum, with tranches 100, 99.9, 99.0 and 90.0 were then applied for the determination of the good versus bad variant annotation profiles. After obtaining the variant annotation profiles, the recalibration was then applied to filter the raw variants. Using VCFTools v.0.1.13 (Danecek *et al*., 2011), SNP trimming and cleaning involved the removal of any SNP with a minor allele frequency (MAF) and “missingness” of < 0.05 and >20%, respectively.

The resultant SNPs were annotated using default parameters for snpEff 4 (Cingolani *et al*., 2012). Ensembl general feature format (GTF, gene sets) information was utilized to build the *P. abies* snpEff database.

### Genetic Structure

A principal component analysis (PCA) was performed on the sampled trees using SNPs derived from the sequence capture data. SNPs with missing values following VQSR were imputed using the nearest neighbour principle in TASSEL (Bradbury *et al*., 2007). This approach was essential considering that PCA demands no missing data points. The covariate matrix derived from the PCA was then displayed by plotting principal component 1 scores against principal component 2 scores in Figure 2. The PCA plot was used to make inference about the population structure. The first two components of the PCA covariate matrix explaining most of the variation were then applied to the AM to account for population structure and correcting for any stratification within the study.

Linkage disequilibrium was calculated using VCFtools v.0.1.13 software using the squared correlation coefficient between genotypes (r2) within scaffolds using the “geno-r2”. The trend-line of LD decay with physical distance was fitted using nonlinear regression (Hill & Weir, 1988) and the regression line was displayed using R (Team, 2015).

### Trait Association Mapping

The LASSO model as described by Li *et al* (2014), Fig. 1c, was applied to all latent traits for the detection of QTLs.

The LASSO model:

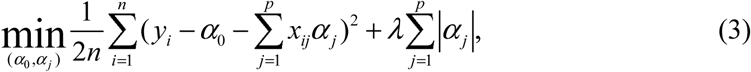
where *y_i_* is the phenotypic value of an individual *i* (*i*=1,…,*n*; *n* is the total number of individuals) for the latent trait *β*_0_, *β*_1_, *β*_2_ or *β*_3_, *α*_0_ is the population mean parameter, *x_ij_* is the genotypic value of individual *i* and marker *j* coded as 0, 1 and 2 for three marker genotypes AA, AB and BB, respectively, *α_j_* is the effect of marker *j* (*i*=1,…,*n*; *n* is the total number of markers), and *λ* (>0) is a shrinkage tuning parameter. A fundamental idea of LASSO is to utilize the penalty function to shrink the SNP effects toward zero, and only keep a small number of important SNPs which are highly associated with the trait in the model.

The stability selection probability (SSP) of each SNP being selected to the model was applied as a way to control the false discovery rate and determine significant SNPs (Gao *et al*., 2014; Li & Sillanpää, 2015). For a marker to be declared significant, a SSP inclusion ratio (Frequency) was used with an inclusion frequency of at least 0.52 for all traits. This frequency inferred that the expected number of falsely selected markers was less than one (1), according to the formula of Buhlmann *et al*, (2014). Population structure was accounted for in all analyses by including the first two principal components based on the genotype data as covariates into the model. An adaptive LASSO approach (Zou et al. 2006) was used to determine the percentage of phenotypic variance (PVE) (*H^2^_QT_*) of all the QTLs (Methods S1). These analysis were all performed in R (Team, 2015), with all the scripts provided in the supplementary material.

### Candidate gene mining

To assess homology of contigs with significant associations, a BLAST search was performed against ConGenIE and public sequence databases, Fig. 1d. After the identification of significant SNPs, the complete *P. abies* contigs that harboured the QTLs were then BLASTed against the ConGenIE database and if no significant hit were detected the whole contig was then extracted. The complete contigs in fasta format were then used to perform a nucleotide BLAST (Blastn) search using the option for only highly similar sequences (megablast) in the National Center for Biotechnology Information (NCBI) nucleotide collection database (https://blast.ncbi.nlm.nih.gov/Blast.cgi?).

## Results

### Norway spruce SNP identification and mapping population structure

All of the 517 Norway spruce mother trees in the study were considered for variant detection and an average of 1.5 million paired end reads were sequenced per individual for the 40019 exome capture probes. This resulted in the identification of 178101 high confidence SNPs. In order to account for effects derived from population stratification we performed a PCA and identified two separate main population groups as well as a number of individuals scattered in between these two main groups. Nevertheless, the differences due to population structure were small with the first two principal components cumulatively explaining only 2.18% of the genetic variation observed (Fig. 3). LD was also determined between all the SNPs, within contigs as well as within significant contigs only and LD decay across physical distance is plotted in Fig 4.

**Fig 3.**
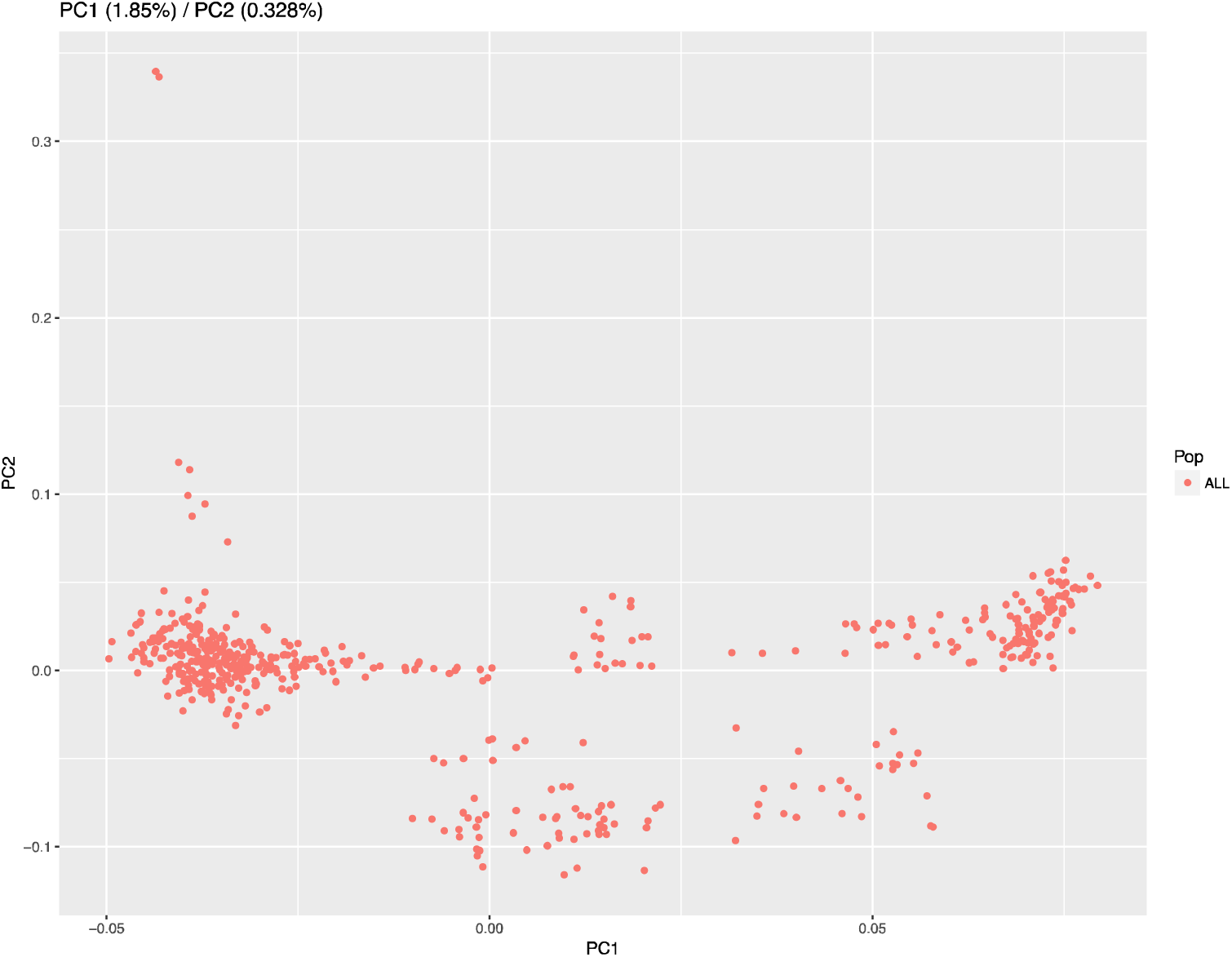
PCA plot of all the 517 mother trees. After VQSR and hard filtering of the SNPs, imputation using the nearest neighbor principle was performed in TASSEL. The PCA indicated a presence of two distinct populations within the 517 mother trees from the Norway spruce breeding program in Sweden. The inferred population structure was used for the correction of stratification within the AM analysis.

**Fig 4.**
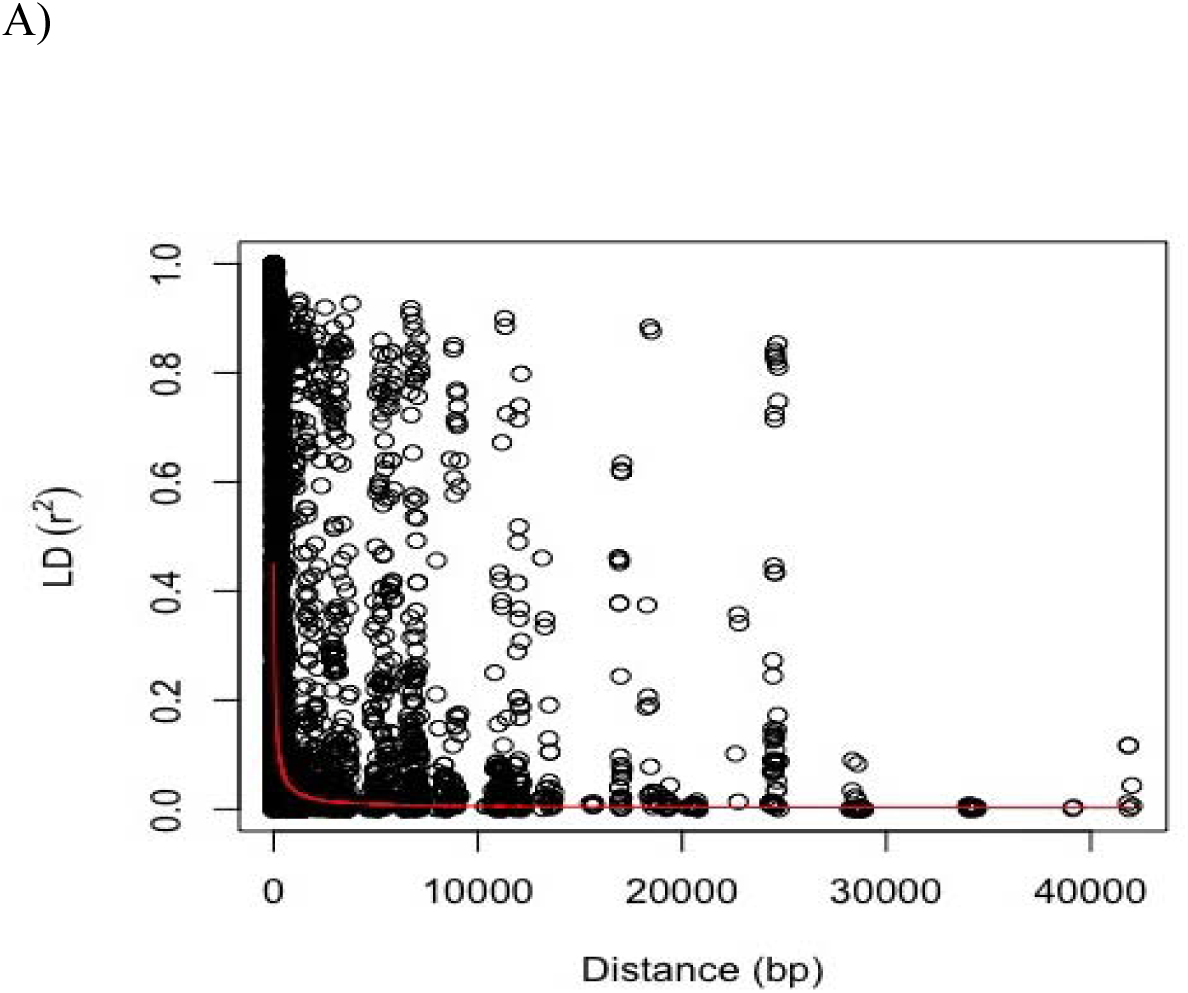
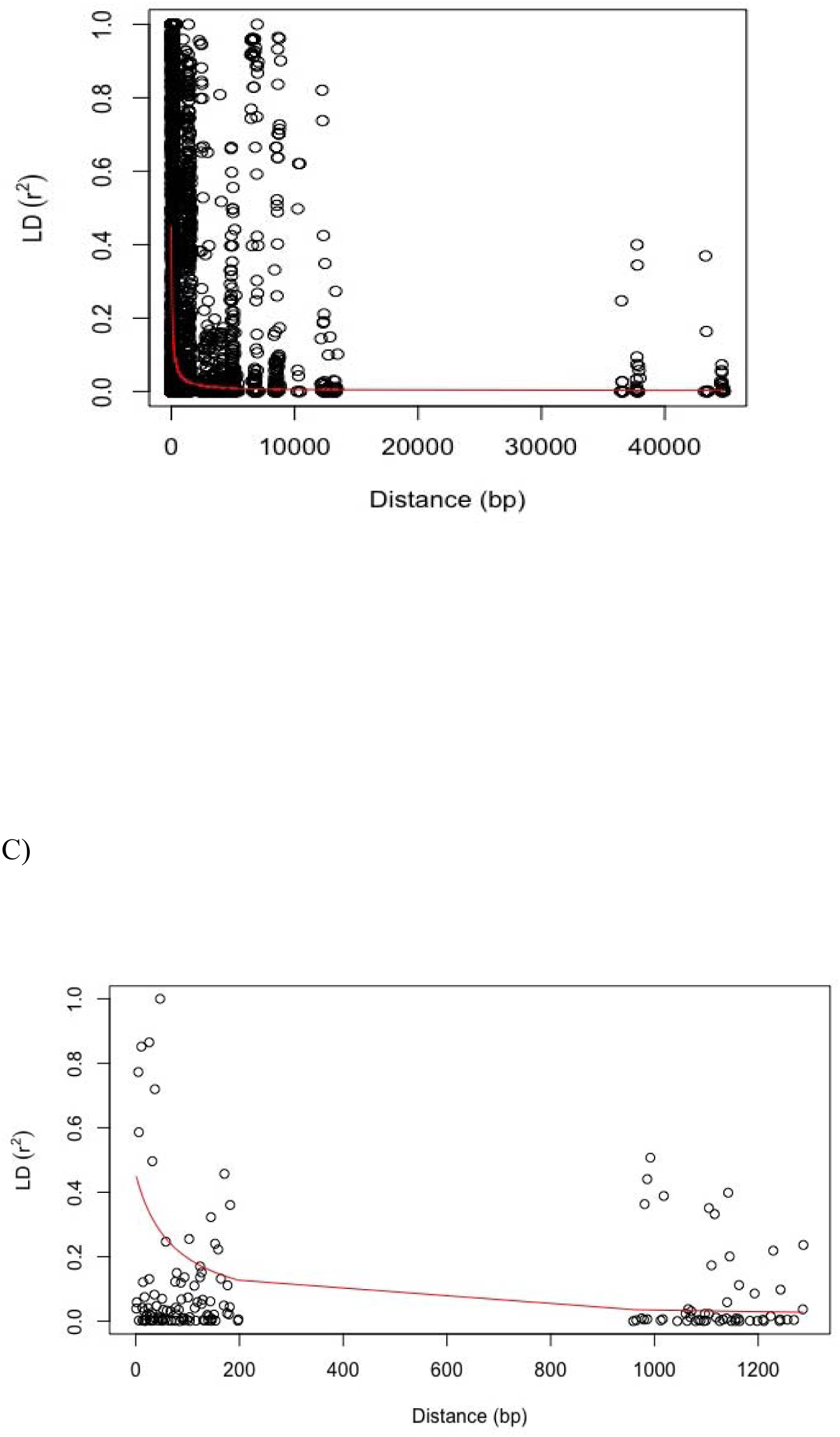
(A) Decay of linkage disequilibrium (LD) across all the tagged genomic sequences, the majority being exoms. Squared coefficients of allele frequency (r^2^) are plotted against distance in base pairs. The fitted curve (red) is representative of the trend of decay from the 178101 SNPs utilised in the association mapping (AM). (B) Decay of LD with distance in base pairs between sites from across 41 contigs with significant associations. (C) Decay of LD across contig MA_96191 that has a significant association for ratio of percentage earlywood vs latewood on which two probes were captured.

### Significant SNPs affecting wood traits

Employing a SSP inclusion frequency of at least 0.52 on the intercept, slope and two knots (K1 and K2) as latent traits, we detected 51 significant QTL across 17 individual traits with the phenotypic variances explained QTL (*H^2^_QTL_*) ranging from 0.01 to 4.93% (Table 2). Several appreciable QTLs were identified with WD and RW having the highest number of associations, at a total of 13 and 14 QTLs, respectively. This was followed by EP/LP-ratio, which had six QTLs. WD, RW and EP/LP were the only three observed traits that have QTLs detected in all four latent traits. For these three phenotypes, the majority of the QTLs were detected when the average ring phenotype was used to derive the latent traits (Table 2). NC associated with one QTL that was detected for the entire ring, whilst six QTLs were identified when EW, TW and LW were analysed separately, with *H^2^_QTL_* ranging from 0.01-4.93% (Table 2).

**Table 2.**
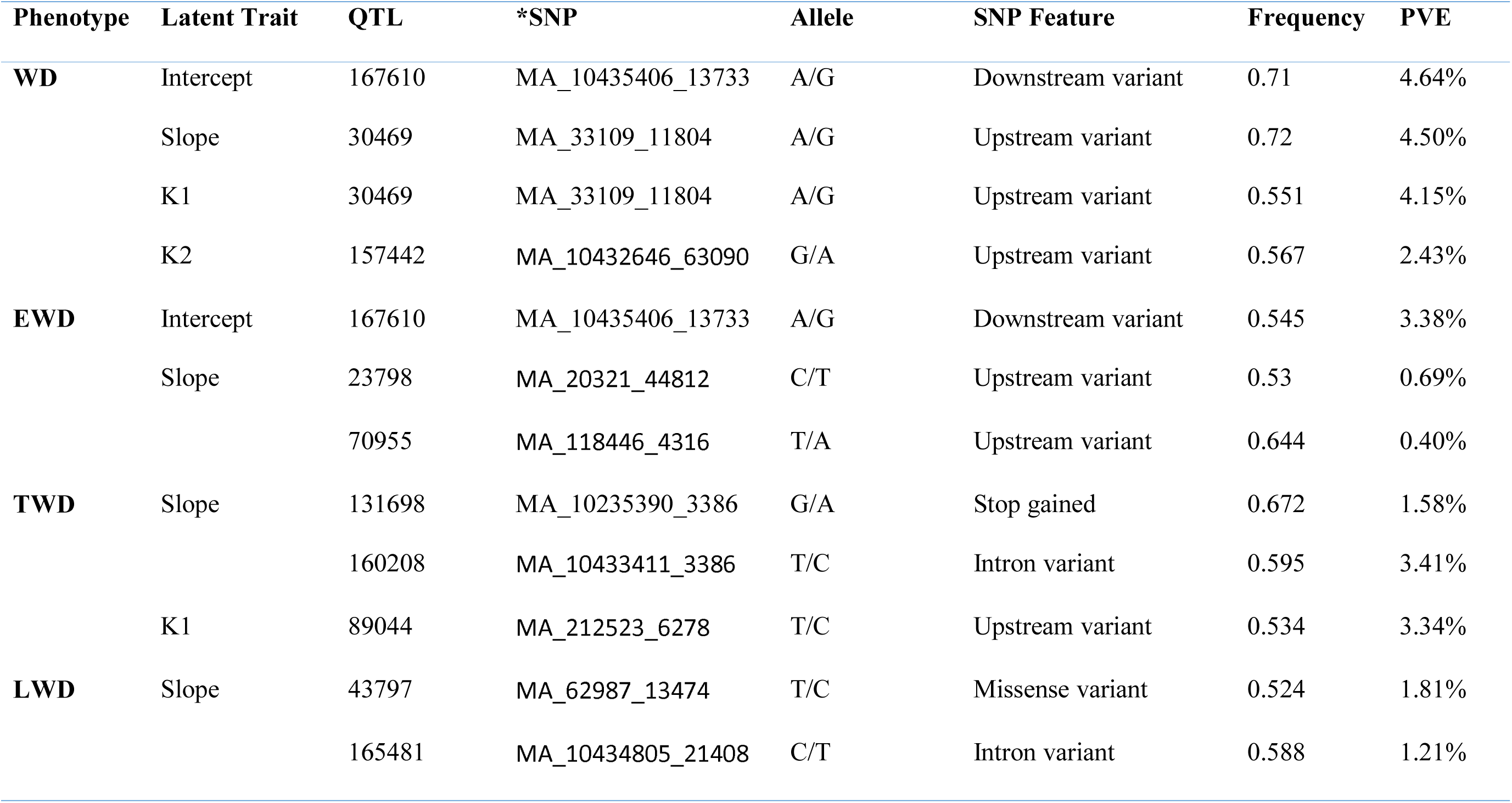
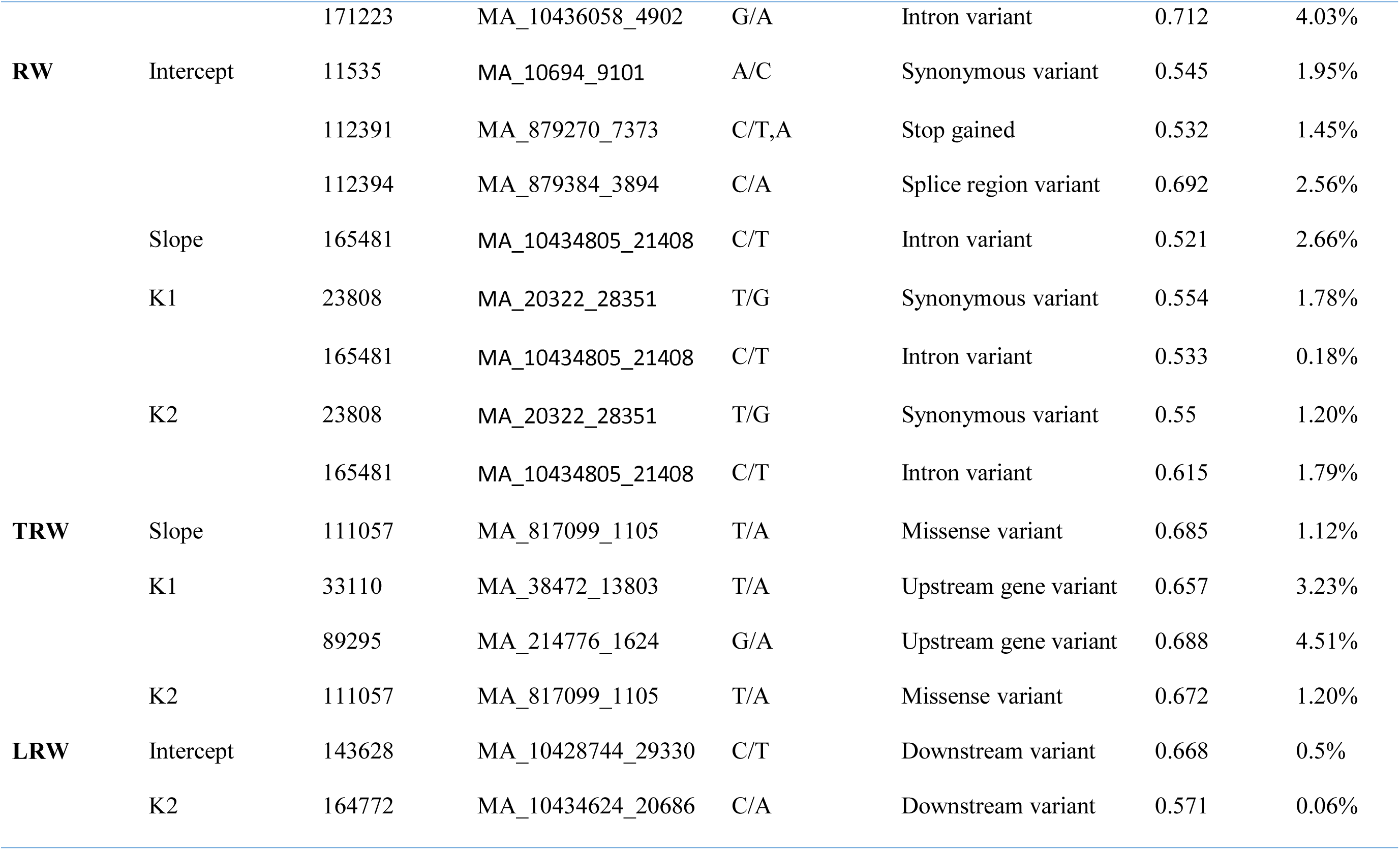
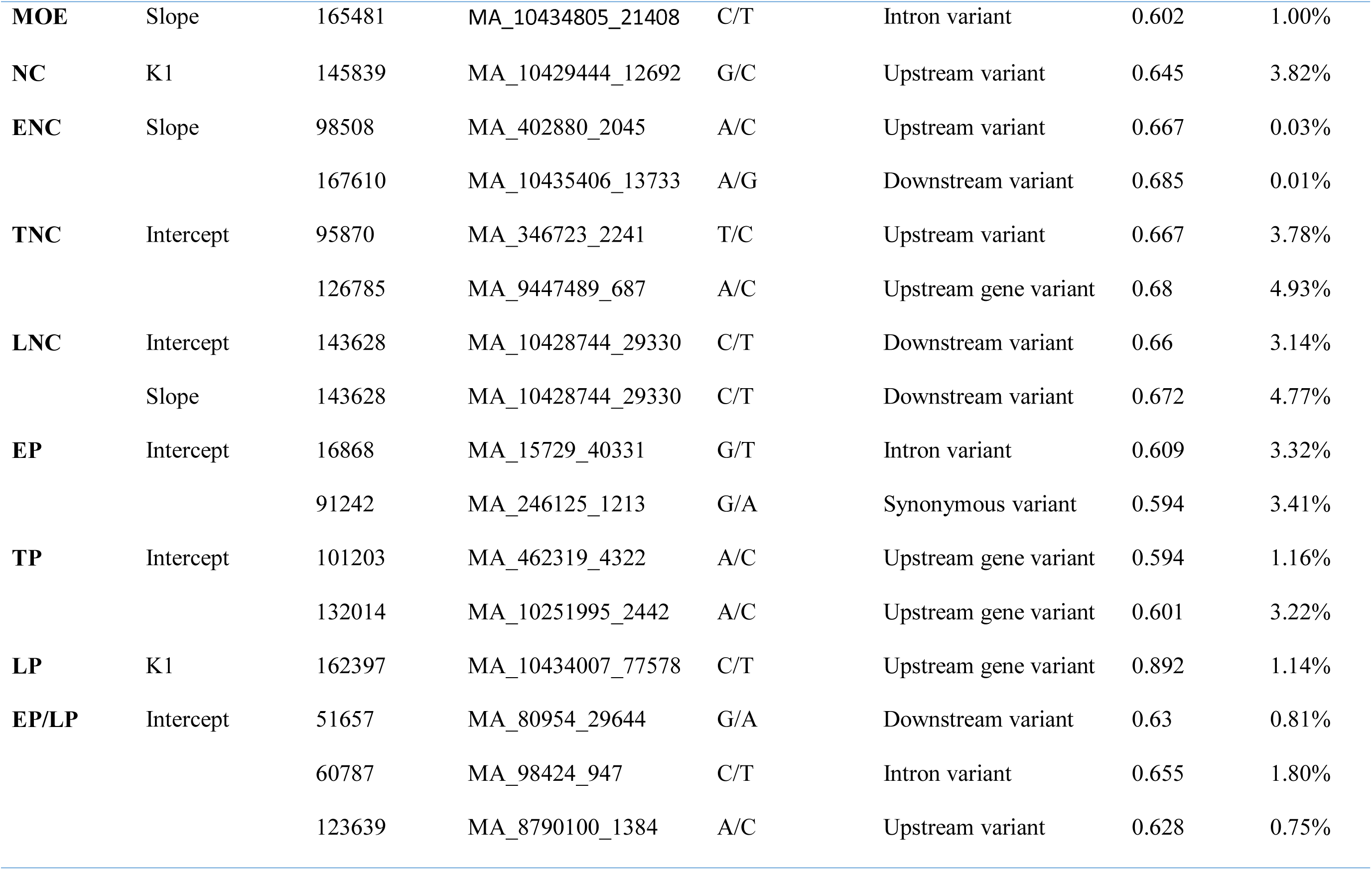
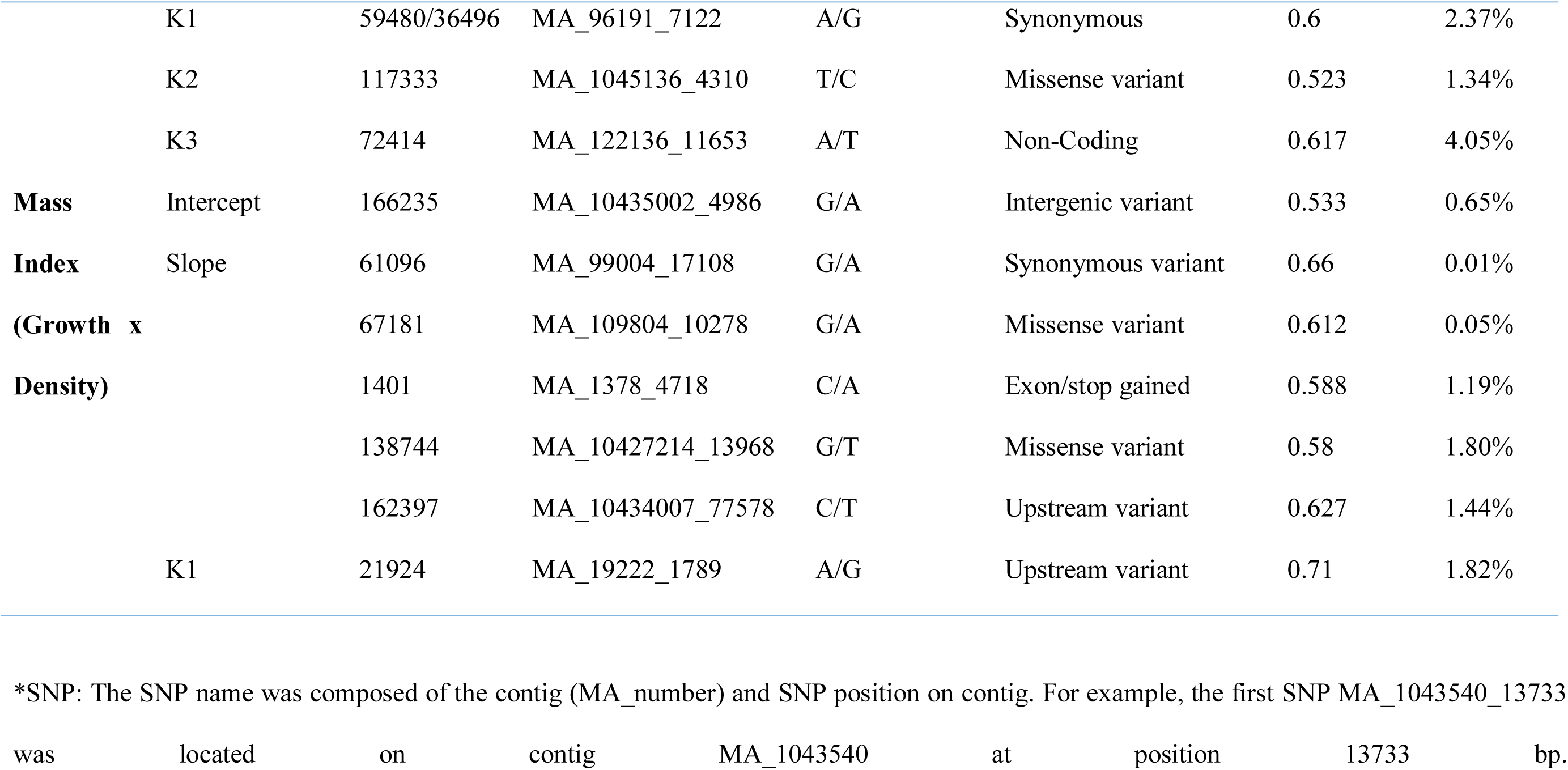
Phenotypes, Latent Traits, SNP, SNP feature, frequency and PVE

Several QTLs shared *within* each trait and *across* traits were observed in the analysis. WD, RW, TRW and LNC had one (30469), two (165481 and 23808), one (111057) and one (143628) QTL shared by two latent traits, respectively. One of the common QTL (30469) for WD had a frequency of 0.72 with an *H^2^_QTL_* of 4.50% for the slope trait, which indicates that it is highly significant for the phenotypes. Common QTLs *within* RW were observed for slope, K1 and K2 latent traits, with moderate frequencies ranging from 0.521 to 0.615 and influenced their respective traits to modest degree (*H^2^_QTL_* in ranges of 0.18-2.66%).

For QTLs common *across* the different latent traits, QTL 165481 was shared between LWD, RW and MOE; this is not surprising because of the close correlation between MOE and wood density, which in turn generally show negative correlation to RW. Intron variant MA_10434805g0010_165481 explained between 0.18-2.66% of the *H^2^_QTL_* observed in the respective traits. The SNP associated with this QTL also had high frequencies of 0.602 and 0.615 in MOE and RW explaining *H^2^_QTL_* of 1.00 and 2.66%, respectively. It was also observed that the SNP MA_10434805g0010_165481 is a common QTL *within* the wood traits related to Width (Table 2). SNP MA_10435406g0010_167610 was shared between WD, EWD and ENC. This SNP was characterized by having high frequencies in WD (0.71) and ENC (0.685), however it had a moderate frequency of 0.545 for EWD. This QTL was detected by the intercept latent trait for WD and EWD, and the slope latent trait in ENC (Table 2), with *H^2^_QTL_* ranging from 0.01-4.64%. The QTL had a high influence on the density related traits as it explained 4.64% (WD) and 3.38% (EWD).

Trees showing a positive correlation between growth and density had seven QTL specific for this observed phenomenon (MI) and had modest influence on the trait (*H^2^_QTL_* in the ranges 0.05-1.82%). Five of the QTL were detected using the slope as the latent trait with high frequencies ranging between 0.58 to 0.66 for SNP 61096 (Table 2).

### Genetic association with phenotypes

Sequence capture and the SNP-trait associations allowed the mining of candidate genes involved in spruce wood formation coupled with the identification of orthologous annotations and descriptions from *Populus* and *Arabidopsis*. This also allowed for the anchoring of significant markers on to the genetic linkage map for Norway spruce Fig 5.

**Fig 5.**
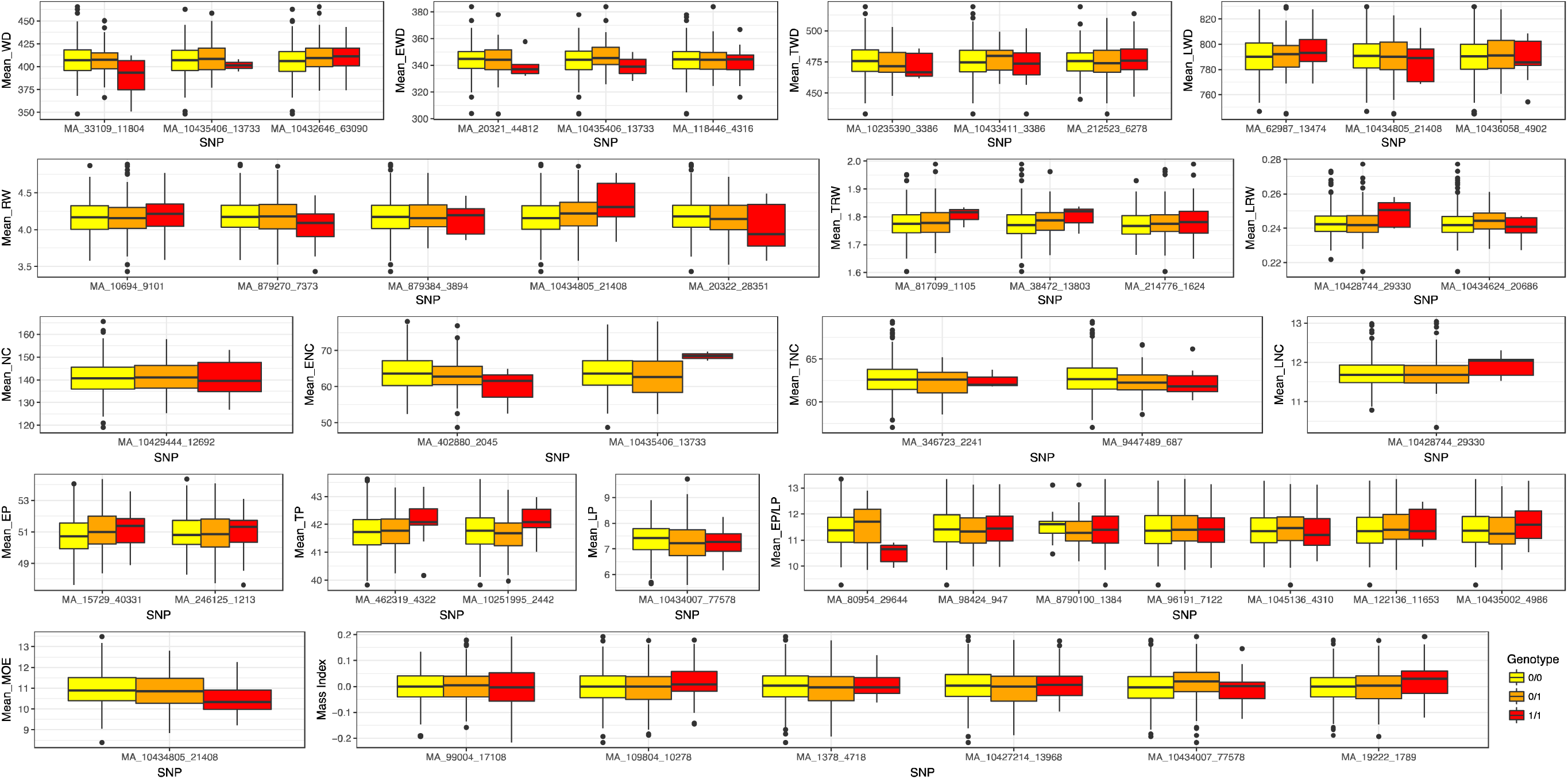
Box plot of the estimated genotypic effect on the phenotypes in the study. The significant SNPs associated and each one of the traits have been correlated to give the impact each genotype has on the average of the overall trait.

**Fig 6.**
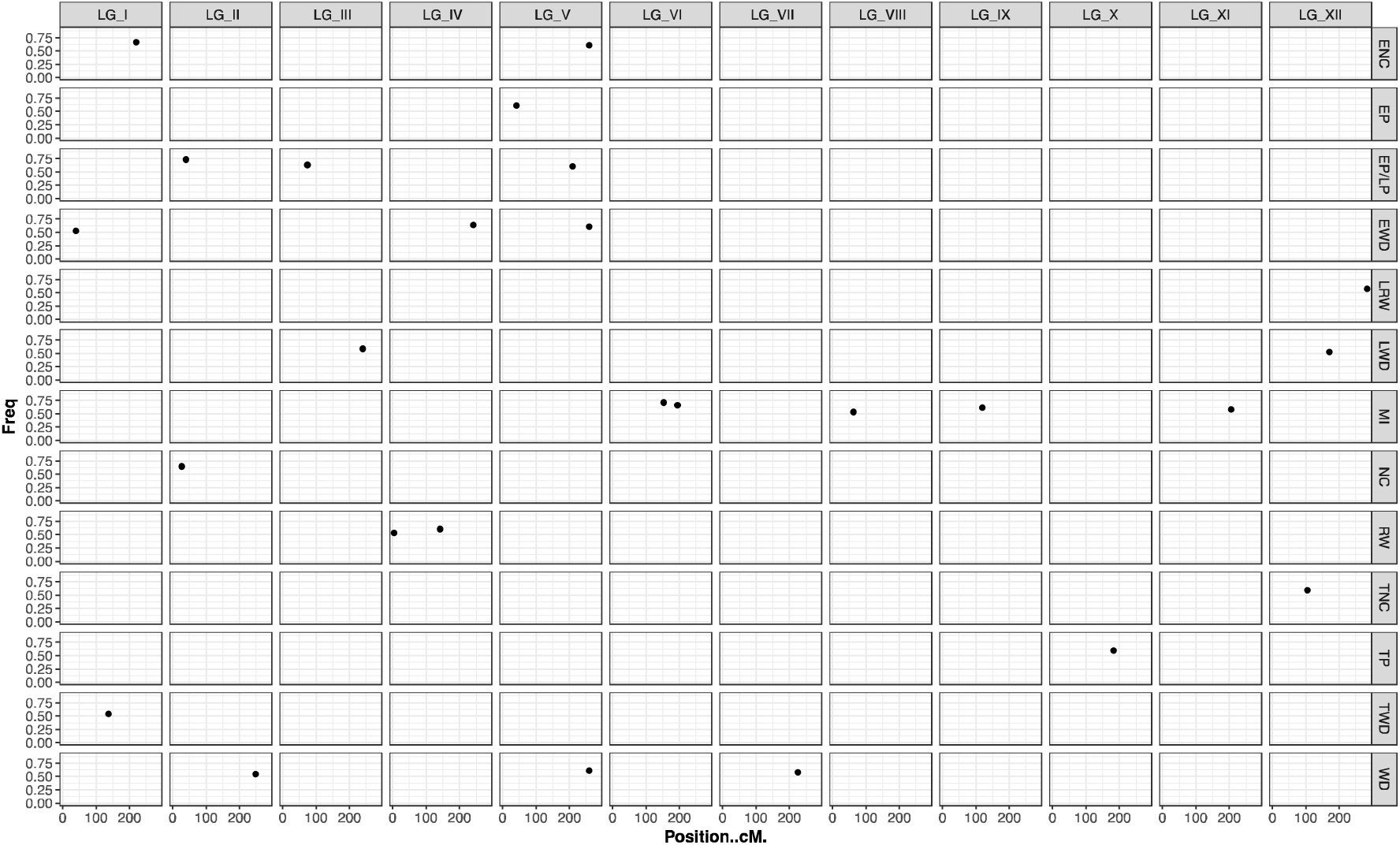
Frequencies of the significant markers selected using the multi-locus LASSO model for whole ring, earlywood and latewood associated with contigs plotted against their locations on a genetic linkage map derived from similar sequence captured probes. Significant associations for the traits were identified on the twelve linkage groups (LG) as follows: [**LG_I**: EWD, TWD and ENC], [**LG_II**: NC, EP/LP and WD], [**LG_III**: EP/LP and RW], [**LG_IV**: RW and EWD], [**LG_V**: EP, EP/LP, ENC, EWD and WD], [**LG**_VI: MI], [**LG_VII**: WD], [**LG_VIII**: MI], [**LG_IX**: MI], [**LG_X**: TP], [**LG_XI**: MI] and [**LG_XII**: TNC, LWD and LRW].

RW, TRW, and LRW were associated with nine gene models. For RW five genes, endoglucanase 11-like, Alpha-dioxygenase 1 (DIOX1), Proliferating cell nuclear antigen (PCNA), B3-DNA-binding and E3 ubiquitin-protein ligase were identified. The SNP MA_879270g0010_112391, a splice region variant, explained 2.56% of the *H^2^_QTL_* and is associated with DIOX1. Marker MA_20322g0010_23808 for RW is associated with the protein domain for a plant specific B3-DNA binding protein, explaining 1.78% variation, with similar orthologous genes in *Arabidposis* and *Populus* (Table S1). The three putative genes associated with TRW are a Serine/threonine-protein kinase, a Homeodomain protein (HB2) and a Senescence-associated protein, and all have high *H^2^_QTL_* ranging from 2.14 to 4.50%. Contig MA_10434624 is homologos to a Pectin esterase and was associated with the downstream variant MA_10434624g0010_164772 for LRW. This may suggest a link between LRW and pectin modification. QTL associated with gene MA_214776g0010 for the TRW may be linked with serine/threonine-protein kinase gene (Os01g0689900), this occurrence of kinase-like related genes was also observed across TRW, NC, EP, EP/LP and EWD (Table S1).

NC, ENC, TNC and LNC are associated with a total of three putative genes and three protein domains. Of the three putative genes, two are associated with serine/kinase activity and one is involved in cysteine and methionine synthesis (Table 1 S1). All the SNPs associated with these traits were either downstream or upstream of coding regions and may thus act as modifiers of gene expression. The SNP MA_402880g0010_98508 (an upstream gene variant) significantly associated with ENC located on gene MA_402880g0010 is homologous to a *Populus* sphingolipid biosynthesis protein. SNP MA_9447489g0010_126785 associated with TNC was located in the gene MA_9447489g0010 which is homologous to a peptidase domain from *Arabidopsis* and showed the highest *H^2^_QTL_* in the dataset (4.93%). This domain is similar to an orthologous zinc carboxypeptidase enzyme of *Oryza sativa* (Zn-dependent exopeptidases superfamily protein) (Table S1).

Wood percentage traits, EP, LP, TP and the ratio of EP/LP had significant associations with ten SNPs. Four of the six significant SNP variants for EP/LP are modifiers with the other two SNPs, being a synonymous (MA_96191g0010_59480) and missense (MA_1045136g0010_117333) variant. The synonymous SNP MA_96191g0010_59480 was associated with the gene model MA_96191g0010, which is homologous to a *P. sitchensis* Glycosyltransferase (GT), similar to UDP-glucosyltransferase 73B2 (AT4G34135) from *Arabidopsis*. Five protein domains were also detected, that were linked to phytochrome kinase substrate 1, TIR/NBS/LRR and zein binding domains (Table S1). The significant SNP MA_15729g0010_16868, an intron variant, that is associated with EP, is located in the gene MA_15729g0010, which is homologous to a DNA-3-methyladenine glycosylase II enzyme. The SNPs identified for TP and LP are all downstream gene variants (Table 2).

WD, EWD, TWD and LWD had a total of 12 significant associations. For the associations with WD we identified the SNP MA_10435406g0010_167610 that is a 3′-gene variant which explained the highest *H^2^_QTL_* observed (4.64%) and is located in a gene that is homologous to a Phosphoadenosine phosphosulfate reductase gene cysH_2. This locus was also detected for EWD and ENC explaining *H^2^_QTL_* of 3.38% and 0.01%, respectively. A missense SNP, MA_33109g0010_30469, was associated with WD and located within the gene MA_33109g0010 homologous to an *Arabidopsis* senescence associated gene 24 (Table S1). The three significant SNPs identified for EWD were all modifiers, upstream and downstream gene variants. Of the three significant SNP associations for TWD, two, SNP MA_10235390_131698 (stop gained) and SNP MA_212523g0010_89044 (upstream gene variant), were identified within genes. The intron variant MA_10433411g0010_160208 associated with TWD and is found in the gene MA_10433411g0010 that is homologous to an *Arabidopsis* Transducin/WD40 repeat-like superfamily protein. Two of the three significant SNPs identified for LWD were intron variants (MA_10434805g0010_165481 and MA_10436058g0010_171223) with the third being a missense variant (MA_62987g0010_43797). The SNP MA_10434805g0010_165481 was found in the gene MA_10434805g0010, which is homologous to an *Arabidopsis* Proliferating Cell Nuclear Antigen Protein (PCNA). This SNP is also associated with RW and explained 1.21% and 2.66% *H^2^_QTL_*, respectively.

The Mass Index trait, that is linked to a positive effect of wood volume growth and increased density (growth x density) yielded a total of seven associated SNPs, with two upstream gene variants, two missense variants one intergenic variant, one stop gained variant and one synonymous nucleotide replacement (Table 2). The slope latent trait had five genes with modest influence on the phenotype ranging from 0.01-1.80%. The genes were homologous to *Arabidopsis* GRAS transcription factor, Aluminium induced protein, Protein virilizer, ARM repeat superfamily protein and an uncharacterized protein. The SNP MA_1378g0010_1401 encodes for a premature stop codon (stop gained, a high impact variant) on gene MA_1378g0010, which is homologous to an *Arabidopsis* protein virilizer involved in mRNA splicing regulation. The gene homologous to the GRAS transcription factor was associated with the SNP MA_99004g0010_61096 (a synonymous variant). The SNP MA_19222g0010_21921, an upstream gene variant, was located in the gene MA_19222g0010 which is homologous to a *Picea sitchensis* ADP (NB-ARC domain) and explained the highest *H^2^_QTL_* of 1.82% (Table S1).

Wood density traits were associated with a total of 12 genes, the largest number of genes identified from the contigs. Percentage of wood was linked to ten putative genes, cell width had nine putative genes and number of cells was associated with six genes. Two genes were shared *across* multiple traits, PCNA was common *across* RW and LWD, and phophoadenosine phosphosulfate reductase was shared *across* WD, EWD and ENC. Genes with the Serine/threonine-protein phosphatase and TIR-NBS-LRR domains were also identified *across* width, wood density and cell percentage traits.

## Discussion

We applied a functional mapping approach in a genome-wide association mapping context and identified 51 significant QTLs that were associated with wood formation in Norway spruce. Previous work utilizing a functional mapping analysis in forest trees have used a limited number of molecular markers (Li *et al*., 2014; Ma *et al*., 2002). Li *et al*., applied this analysis in a bi-parental Scots pine (*P. sylvestris* L.) cross using 319 markers. Hence, our work represents a major advance in that we have been able to apply this approach at a genome-wide scale (178101 SNPs) on unrelated mother trees, with a dynamic functional trait dataset comprising 15-time points/annual growth rings (*i.e*., cambial age). Latent traits represent significant time points in the trait development allowing us to detect putative genes at these critical junctures in wood formation.

The number of detected QTLs is relatively small compared with several recent studies in *Populus* (Evans *et al*., 2014; McKown *et al*., 2014; Porth *et al*., 2013). The sample size, number of SNPs used and the stringency with which we accepted significant SNPs contributed to the modest number of QTL. Previous functional mapping studies, (Li *et al*., 2014) involving SNPs in conifers have used two levels of evaluating QTLs, whereby they have suggestive and significant QTL. In our study, we only reported significant QTL. As indicated in Hall *et al* (2016), there should be hundreds to thousands of QTL of moderate to very small effect related to growth and wood quality traits in trees. Hence, a large population and accurate phenotyping are required for a reliable identification of most QTLs (Korte & Farlow, 2013). However, the sample size of our study allowed the detection of the largest/most significant QTL. The study identified significant associations explaining relatively small proportions of the phenotypic variance being observed, ranging from 0.01-4.93%. This is in line with other studies of QTL for wood traits (González-Martínez *et al*., 2007; Porth *et al*., 2013).

### Genetic associations with potential to improve wood properties

With all the SNPs, having been derived from known genomic positions, it was possible to identify genes linked to the associated QTLs and infer their potential function in wood formation.

The gene MA_10694g0010 is homologous to an enzyme involved in cell wall biosynthesis, endoglucanase 11-like, and was associated with RW (intercept latent) (Table S1). The association of this gene with the RW intercept implies that the gene influences the trait throughout the growth period. This enzyme is a vital component of xylogenesis and is involved in the active digestion of the primary cell wall (Goulao *et al*., 2011). The endoglucanase 11-like, was associated with a synonymous SNP MA_10694g0010_11535 for (RW) suggesting an involvement in cell expansion and cell wall loosening during wood formation. Endoglucanases have been proposed as enzymes involved in controlling cell wall loosening (Cosgrove, 2005). Endoglucanase 11-like gene is part of the endo-1 family in which the eno-1-4-β-glucanase *Korrigan* gene belongs. Characterisation of the *Korrigan* gene in *P. glauca* has identified it to be functionally conserved and essential for cellulose synthesis (Maloney *et al*., 2012). Hence, MA_10694g0010 is a candidate for the remodelling of cell walls that affects the mechanical and growth properties of wood cells, and consequently annual ring width.

The synonymous SNP MA_20322g0010_23808 is associated with RW located on gene MA_20322g0010 which is homologous with a plant specific B3-DNA binding protein domain, that is shared among various plant-specific transcription factors. This includes transcription factors involved in auxin and abscisic acid responsive transcription (Yamasaki *et al*., 2004). Auxin is one of the central phytohormones in the control of plant growth and development (Abel & Theologis, 1996), and also known to be involved in cell wall loosening and elongation (Cosgrove, 2016). This suggests a possible functional role for MA_20322g0010 in influencing RW.

An intron variant located in the MA_10434805g0020 gene, which is homologous to PCNA was detected *across* several phenotypes (LWD, RW and MOE) associated with the slope latent trait (Table 2). The detection of this gene *across* these phenotypes using the slope latent trait implies that the gene affects the rate of change of these phenotypes. Thus, this would be a good gene to target for further studies. PCNA proteins function as integral enzymes in the regulatory pathways of cell cycle regulation and DNA metabolism (Maga & Hübscher, 2003). PCNA has been associated with chromatin remodelling, DNA repair, sister-chromatid cohesion and cell cycle control, which are all vital processes in plant growth (Strzalka & Ziemienowicz, 2010), but it has not been previously associated with wood formation traits.

In our study we detected a significant downstream SNP (MA_10434624g0010_164772) associated with LRW on gene MA_10434624g0020, homologous to pectinmethylesterases (PMEs), which are cell wall associated enzymes responsible for demethylation of polygalacturonans (Phan *et al*., 2007). This enzyme has been shown to be linked with many developmental processes in plants, such as, cellular adhesion and stem elongation (Micheli, 2001). An association study in White spruce identified a significant nonsynonymous SNP coding for cysteine associated with earlywood and total wood cell wall thickness associated with pectinmethylesterase (Beaulieu *et al*., 2011). Our study identified a PME SNP association in the latewood stage, supporting the importance of PMEs in wood cell development.

A SNP (MA_10435406g0010_167910) downstream on gene MA_10435406g0010 was detected *across* the traits ENC, WD and EWD. The association of this gene with the WD and EWD intercept implies that it an impact on the overall development of density throughout the growth period. Since density is correlated with number of cells, this association with the slope latent trait of ENC means the gene influences its rate of change. The gene is homologous to Phosphoadenosine phosphosulfate reductase (PAPS), which plays a central role in the reduction of sulphur in plants. An analysis of PAPS enzymes in *Arabidopsis* (Klein & Papenbrock, 2004) and *Populus* (Kopriva *et al*., 2004) revealed that enzymes involved in sulphate-conjugation, play an important role in plant growth and development (Klein & Papenbrock, 2004). Reduced sulphur is utilized by the sulphate assimilation pathway for the synthesis of essential amino acids cysteine and methionine (Kopriva & Koprivova, 2004). Methione acts as a methyl donor in both lignin, hemicellulose and pectin biosynthesis providing a possible mechanism of how PAPS could influence wood density and number of cells.

When analyzing QTLs detected for traits linked to the percentage of cells (EP, LP and EP/LP) we identified three putative candidate genes, DNA-3-methyladenine glycosylase II enzyme, phytochrome kinase substrate 1 and glycosyltransferase. DNA-3-methyladenine glycosylase II enzyme is responsible for carrying out base excision repairs (BER) in the genome in order to maintain genomic integrity. This enzyme has the ability to initiate a broad substrate recognition and provides a wide resistance to DNA damaging agents (Wyatt *et al*., 1999). This DNA repair capacity can be expected to be essential for the process of cell propagation and growth.

A synonymous SNP (MA_96191_59480) within the gene MA_96191g0010, which is homologous to Glucosyltransferase in *P. sitchensis* was associated with EP/LP. Glycosyl transferases operate by facilitating the catalytic sequential transfer of sugars from activated donors to acceptor molecules that form region and stereospecific glycosidic linkages (Lairson *et al*., 2008). The Arabidopsis ortholog (UDP-glucosyltransferase 73B2) encodes for a putative flavonol 7-O-glucosyltransferase involved in stress responses. In our study, this significant association was associated with EP/LP, however a nonsynonymous variant in a gene coding for a Glycosyl transferase in *Populus* was associated with fibre development and elongation (Porth *et al*., 2013). Therefore, gene MA_96191g0010 is a novel candidate for further investigation of how flavonol metabolism may influence the proportion of early and late wood in Norway spruce.

Two genes concerning wood formation, PAPS and PCNA, were also detected across related traits density, growth number of cells and MOE. Significant SNP (MA_10435406_167610) in the PAPS reductase gene is common *across* ENC, WD and EWD, with SNP MA_10434805_165481 located in an intron for a gene encoding for PCNA protein being detected *across* WD, RW and MOE (Table S1). The presence of these common QTL suggests that these traits might be under the control of the same genes or genetic pathways. Chen et al (2014) reported a significant positive genetic correlation between wood density and MOE, which increased with tree age. However, wood volume growth and density have a negative correlation (Chen *et al*., 2014), with our study being the first to detect QTLs for trees exhibiting a positive correlation for this phenomenon (MI). The common QTL observed *across* WD, EWD and ENC indicates that the number of cells during the juvenile wood development stages has a significant impact on the overall density. The seasonal changes in EWD to LWD has been speculated to be due to a change in auxin levels leading to the initiation of wall-thickening phase, which has a direct impact on the wood quality traits such as MOE. This phase coincides with the cessation of height growth and where available resources are used for cell-wall thickening (Sewell *et al*., 2000), which may explain the common QTL between LWD, RW and MOE, as part of the same feedback loop mechanism.

We identified two associations to homologous genes related to nucleic acid repair functions, DICER-LIKE3 (DCL3) and DNA mismatch repair protein (MSH5), which are concerned with RNA processing as well as DNA repair, respectively. These genes are involved in ensuring the fidelity of DNA replication and to preserve genomic integrity (Hsieh & Yamane, 2008). These genes are possibly associated with cambial cell division and endo-reduplication during wood formation and can conceivably have effects on wood density.

An association for TWD with a SNP located upstream of gene MA_212523g0010, is homologous to Kinesin-related protein 13 (gene-L484_021891). Kinesin-related proteins are known to be involved in secondary wall deposition, which can impact wood density (Zhong *et al*., 2002), cell wall strength and oriented deposition of cellulose microfibrils.

Several receptor-like Kinases (TIR/NBS/LRR and Serine/threonine-protein phosphatase) homologs were identified *across* traits (TRW, NC, EP, EP/LP and EWD) (Table S1). These protein domains control a large range of processes including hormone perception and plant development. Approximately 2.5% of the annotated genes in *Arabidopsis* genome are RLK homologs (Shiu & Bleecker, 2001), where they among other functions play an important role in the differentiation and separation of xylem and phloem cells (Fisher & Turner, 2007). Similar to our study a synonymous SNP in a RLK gene was associated with early wood proportion (EP) in White spruce (Beaulieu *et al*., 2011), hence RLKs seem to be involved in modifying a number of different wood properties from density to cell identity and number.

Norway spruce trees that possess the ability of fast growth and high wood density are very rare, but such trees and associated SNPs were discovered in our study. Trees combining these traits are of high interest to forest industries and owners, and thus also in focus for breeders. Of the seven genes significantly linked to this phenomenon of particular interest was a synonymous SNP on MA_99004g0100 gene homologous to a transcription factor from the GRAS family (Table S1). GRAS is an important class of plant-specific proteins derived from three members: <u>G</u>IBBERELLIC-ACID INSENSITIVE (GAI), <u>R</u>EPRESSOR of GAI (RGA) <u>a</u>nd <u>S</u>CARECROW (SCR) (GRAS) (Hirsch & Oldroyd, 2009). GRAS genes are known to be involved in the regulation of plant development through the regulation of gibberellic acid (GA) and light signalling (Cenci & Rouard, 2017; Hirsch & Oldroyd, 2009). Furthermore GA signalling has also been shown to stimulate wood formation in *Populus* (Mauriat & Moritz, 2009). Thus, the GRAS transcription factor identified here and the other six genes positively associated with MI provide interesting genetic markers and tools to understand this phenomenon.

## Conclusion

This work has dissected the genetic basis of wood properties in Norway spruce with use of functional association mapping. In total, we identified 51 Significant QTLs for wood properties and mining of candidate genes located in the vicinity of significant QTLs identified genes that could be directly or indirectly responsible for variations in the observed traits. Significant novelty in our results is provided by the identification of QTLs associated to both high wood density and fast growth, thus larger biomass. These genes are candidates for further functional verification in Norway spruce.

## Acknowledgements

We acknowledge the support of the Bio4Energy research organization for wood property analyses and evaluations. All genetic data was obtained through funding from the Knut and Alice Wallenberg foundation. SSF project support of continuing this work. JB is supported though a postdoc position funded by the Kempe foundation.

